# *zmiz1a* zebrafish mutants have defective erythropoiesis, altered expression of autophagy genes, and a deficient response to vitamin D

**DOI:** 10.1101/2021.05.14.444216

**Authors:** Francisco Castillo-Castellanos, Laura Ramírez, Hilda Lomelí

## Abstract

ZMIZ1 is a transcriptional coactivator that is related to members of the protein inhibitor of activated STAT (PIAS) family. ZMIZ1 regulates the activity of various transcription factors including the androgen receptor, p53, and Smad3. ZMIZ1 also interacts with Notch1 and selectively regulates Notch1 target genes relevant for T cell development and leukemogenesis in mammals. Human ZMIZ1 is additionally characterized as a latitude-dependent autoimmune disease (LDAD) risk gene, as it is responsive to vitamin D and has been associated with at least eleven blood cell traits.

To address the function of ZMIZ1 in fish, we introduced CRISPR/Cas9 mutations in the *zmiz1a* gene in zebrafish. We observed that inactivation of *zmiz1a* in developing zebrafish larvae results in lethality at 15 dpf and delayed erythroid maturation. Differential gene expression analysis indicated that 15 dpf *zmiz1a*-null larvae had altered expression of autophagy genes, and erythrocytes that lacked Zmiz1a function exhibited an accumulation of mitochondrial DNA. Furthermore, we observed that autophagy gene expression was dysregulated at earlier stages of development, which suggests the involvement of Zmiz1a in the regulation of autophagy genes beyond the process of red blood cell differentiation. Finally, we showed that the loss of Zmiz1a decreased the capacity of the embryos to respond to vitamin D, indicating additional participation of Zmiz1a as a mediator of vitamin D activity.

## 1. Introduction

Hematopoiesis in zebrafish is a highly conserved developmental process. Both the morphology of hematopoietic cell types and the genetic mechanisms involved in the differentiation of blood cells share striking similarities with those processes described in the mammalian models. Similar to higher vertebrates, zebrafish undergo two waves of hematopoiesis: primitive and definitive [1-3]. Primitive hematopoiesis initiates in embryos at approximately 12 hours postfertilization (hpf) with the detection of hemangioblasts that are positive for *gata1*. Hemangioblasts are progenitor cells derived from the ventral mesoderm that arrange in two parallel stripes along the posterior trunk. Later, these cells migrate to the posterior part of the intermediate cell mass (ICM), a tissue that extends along the trunk between the notochord and the yolk sac, where they develop into erythroid cells. By 24 hpf, primitive erythroblasts enter the circulation and account for the blood function for at least 4-5 days, when erythrocytes derived from the definitive wave incorporate into the blood [1, 4-6]. Definitive hematopoiesis is initiated in 26-28 hpf embryos in the region located between the axial vein and the dorsal aorta. Hematopoietic self-renewing progenitor cells (HSPCs) are generated by transdifferentiation from endothelial cells of the ventral aorta in a process that is known as endothelial-to-hematopoietic transition [7-9]. HSPCs generate pluripotent hematopoietic stem cells (HSCs), which have the potential to differentiate into all blood cell types and are recognized by the expression of *runx1* and *cmyb*. As they originate, HSCs enter the circulation and start a migration that sequentially seeds the locations where different lineages will be produced [3, 5, 6]. By 48 hpf, HSCs reach the caudal hematopoietic tissue through the axial vein. Six hours later, they arrive in the thymus, which is an exclusive site for the production of T lymphocytes, and by 4 dpf, HSCs colonize the kidney marrow, which becomes the main organ for hematopoiesis in adults throughout the lifespan.

Pluripotent HSCs give rise to progenitors of different lineages, including erythroid-committed progenitors. In humans, erythroid progenitors are classified according to their *in vitro* colony-forming properties into burst-forming unit-erythroid (BFU-E) cells and colony-forming unit-erythroid (CFU-E) cells. CFU-E cells undergo three to five divisions during their differentiation and ultimately produce mature erythrocytes [10]. In zebrafish, erythrocyte maturation starts with the generation of proerythroblasts, cells with a strong basophilic cytoplasm and granular nuclei. Maturation involves the progressive acidification of the cytoplasm, the condensation of nuclear chromatin, the loss of mitochondria and a reduction in cell diameter. Mature fish erythrocytes are elongated cells that conserve a small nucleus [11]. The key factor that controls the gene transcription programs required for the terminal differentiation of erythroid cells is Gata-1 [12].

Human ZMIZ1 protein was initially identified as an androgen receptor (AR) coactivator [13]. Analysis of its sequence revealed the presence of several important functional domains, including a tetratricopeptide repeat (TPR) at the N-terminus extremity, an alanine-rich motif predicted to be intrinsically disordered, and two proline-rich domains, one of them within the C-terminus, containing a strong transactivation activity (TAD) and a SP-RING/Miz domain, which is highly conserved in members of the PIAS family and confers SUMO-E3-ligase activity to the PIAS proteins [14, 15].

In addition to coactivating the transcription mediated by the AR, Zmiz1 is known to coactivate p53, Smad3 and Notch signaling [16-18]. Importantly, it has been shown that during T cell development in mice, Zmiz1 promotes normal and leukemic pre-T-cell proliferation. For this function, Zmiz1 interacts directly with Notch1 through its N-terminal domain (the TPR) and regulates more than 40% of Notch-induced target genes [18]. Consistently, mutagenesis screens implicated ZMIZ1 as a NOTCH1 collaborator in human T-cell acute lymphoblastic leukemia [19]. Further associations of ZMIZ1 with severe human diseases have been discovered in several GWAS and case studies. In particular, hZMIZ1 is a risk factor for several autoimmune conditions. Among the traits that have been associated with ZMIZ1 are multiple sclerosis (MS) [20-22], Crohn’
ss disease [23-25], inflammatory bowel disease [26, 27], celiac disease [28, 29], psoriasis [25], vitiligo [30, 31] and dysmenorrhea [32]. Interestingly, the regulation of ZMIZ1 by vitamin D appears to contribute to autoimmunity. Additional GWAS studies found ZMIZ1 to be a risk allele for type 2 diabetes [33, 34], and in other genomic screens, ZMIZ1 has been identified as an oncogene implicated in breast [35, 36], epithelial and colorectal cancer [37-39]. Finally, in a recent study, Carapito *et al*. found that ZMIZ1 variants with mutations that affect key protein regions cause a syndromic neurodevelopmental disorder [15]. Notably, in these studies, ZMIZ1 expression was found downregulated in autoimmune disorders, whereas it was found upregulated in cases of cancer, which highlights the importance of the precise regulation of *ZMIZ1* expression in human cells.

The multiple phenotypes produced by abnormalities in ZMIZ1 expression are not surprising considering the various functional domains of ZMIZ1, which entail a diversity of potential interactions with a broad range of transcription factors. Therefore, an important role of ZMIZ1 during embryonic development was anticipated. On this subject, it was shown that mice homozygous for a mutation in *Zmiz1* die at mid-gestation due to yolk sac vascular remodeling failure and abnormal embryonic vascular development [40].

In this study, we used zebrafish to analyze the roles of Zmiz1a during embryonic development. Zebrafish present advantages for studies involving cardiovascular or blood phenotypes since their oxygen homeostasis is independent of blood circulation, and thus, they can form organs in the absence of a functional cardiovascular system. We obtained CRISPR/Cas9 *zmiz1a* mutant fish and found that in the absence of Zmiz1a, most of the larvae die between 15-20 dpf, whereas approximately 30% survive to adulthood. Homozygous mutant larvae displayed abnormalities in definitive erythropoiesis, which consists of delayed erythroid maturation observed from 11 dpf, reaching its maximum penetrance at 15 dpf, when embryos started to die. Transcriptomic analysis of the mutants indicated that the gene groups with the highest enrichment of differentially expressed genes (DEGs) were those associated with autophagy, mitochondrion organization and the biosynthesis and metabolism of sterols and steroids. We show that circulating erythropoietic cells in the *zmiz1a* mutant contain increased copy numbers of mitochondrial DNA, suggesting a defective elimination of mitochondria. Additionally, we found that autophagy gene expression is altered in the mutant embryos starting in the early stages, which suggests a role of Zmiz1a in the regulation of autophagy genes beyond the process of red blood cell differentiation. Finally, we studied the vitamin D response in *zmiz1a* mutants and found a reduced capacity of the *zmiz1a*^*-/-*^ embryos to respond to calcitriol treatment. This last finding is consistent with the previous findings that indicate a relevant participation of Zmiz1a in the immune response.

## 2. Materials and methods

### 2.1. Fish maintenance and strains

A zebrafish (*Danio rerio*) AB-TU-WIK hybrid line was used. The embryos were obtained by natural crosses and raised at 28 °C based on standard procedures [41]. The staging was performed according to the Kimmel system [42]. Zebrafish transgenic lines *Tg(mpx:gfp)*^*i114*^ and *Tg(mpeg1:EGFP)*^*gl22*^ used respectively for neutrophile and macrophage lineage tracking (Renshaw y Ellet), were generously donated by Dr. Herman Spaink. Zebrafish were handled in compliance with local animal welfare regulations and all experimental protocols were approved by the Comité de ética (Instituto de Biotecnología, UNAM).

### 2.2. In situ hybridization

The RNA *in situ* hybridization using DIG-labeled antisense RNA probes was performed according to reported standard protocols [43]. The plasmids used for *in situ* probe synthesis were previously described and generously donated as follows: *gata1* by Leonard Zon [44], *runx* by Katherine Crosier [45] and *rag1* by Catherine Willett [46]. For synthesis of the *zmiz1a in situ* probe the *zmiz1a* fragment from 1078 to 1747 from the cDNA sequence was cloned and transcribed.

### 2.3. CRISPR/Cas9-mediated mutations and genotyping

CRISPR/Cas9 target sites were designed using an online tool ZiFiT Targeter software (http://zifit.partners.org/ZiFiT). The *zmiz1a* genomic target sequence is 5′GGACCTCGGCTACCGGCTTCTGG3′, located at exon 3. The primers 5′AAACGAAGCCGGTAGCCGAGGT3′ and 5′TAGGACCTCGGCTACCGGCTTC3′ were annealed and cloned into the pDR274 plasmid [47]. sgRNA was synthesized using T7 RNA polymerase (Roche). AmpliCap SP6 High Yield Message (CellScript) was used for the Cas9 mRNA synthesis using the pCS2-nls-zCas9-nls plasmid [47]. One-cell stage embryos were injected directly into the cell with ∼10 ng/µl of sgRNA, and ∼40 ng/µl of Cas9 mRNA diluted in 100 mM KCl. Embryos injected only with Cas9 were used as controls.

The targeted genomic locus was amplified with Phusion High-Fidelity DNA Polymerase (Thermo) from single embryos or larvae using primers that anneal to sites 50 and 385 bp upstream and downstream from the target site respectively (see supplementary table 1). The PCR product was cloned into a pJet1.2 plasmid (Thermo Scientific) for sequencing. Primers flanking the target site were used to amplify a 252 bp fragment for genotyping (supplementary table 1).

### 2.4. Survival Assay

Crosses from *zmiz1a+/-* fishes were raised at 28 °C with a density of a hundred larvae per 300 ml. From each cross 20 larvae were sampled at each time point 1, 9, 11, 13, 15, 17 & 19 dpf for genotyping. Surviving larvae were calculated based on the distribution of genotypes. The survival difference and statistical significance were determined by a log-rank test.

### 2.5. Histology

After in situ hybridization, embryos were fixed overnight at 4 °C in Bouin’
ss solution and dehydrated by a series of graded ethanol. Samples were embedded with paraffin after xylene. Tissue sections of 10 µm were cut and stained in hematoxylin-eosin based on a reported protocol [48]. Subsequently, sections were mounted and photographed on a Leica DMLB microscope equipped with an AxioCam MR5 (Zeiss) camera.

### 2.6. Microscopy and analysis

For live imaging, embryos were anesthetized using 0.016% tricaine (Sigma). Both live and fixed embryos were mounted in 0.6% low-melting agarose. Fluorescent and brightfield image acquisition was performed using a Leica DMLB 100S stereoscope. For erythrocytes slides image acquisition was performed on a Leica MZ12S microscope under Normarski illumination. Images were processed with Infocus and ImageJ software. Images were adjusted for brightness and contrast using Image J.

### 2.7. Embryonic Blood Collection and Morphometry

Embryonic zebrafish erythrocytes at indicated times after fertilization were collected by transecting tails of 10–12 embryos in 20 µL of PBS. Larvae were anesthetized and left to bleed for 5 minutes before collecting the bodies for genotyping. Slides were prepared and stained with Wright/Giemsa stain (Hycel) according to the manufacturer’s instructions. Embryonic erythrocytes were stained with hemoglobin-specific O-dianisidine (Sigma-Aldrich) staining as described previously [49]. Nuclear and cytoplasmic areas of 60–80 randomly selected erythrocytes were measured by using Image J software, and the N:C area ratio was calculated.

### 2.8. RNA-sequencing and data analysis

Pools of twenty 15 dpf-embryo (*zmiz1a*^*+/-*^ and *zmiz1a*^*-/-*^) were used for total RNA isolation with RNAeasy columns (Qiagen) from five independent crosses. Each RNA sample was analyzed for quantity and purity with Agilent 2100 Bioanalyzer (Agilent Technologies, CA, USA). RNA samples were used for RNA sequencing (4 control *zmiz1a*^*+/-*^ embryos and 3 *zmiz1a*^*-/-*^ mutant embryos). RNA sequencing and data analysis were performed by Novogene Co., Ltd. According to their protocols, sequencing libraries were generated using NEBNext®Ultra™RNA Library Prep Kit for Illumina® (NEB, USA) following the manufacturer’s recommendations, and index codes were added to attribute sequences to each sample. Library preparations were sequenced on an Illumina platform and 125 bp/150 bp paired-end reads were generated. Raw reads were filtered into clean reads aligned to the reference sequences. The alignment data was utilized to calculate the distribution of reads on reference genes and mapping ratio. The fragments per kilobase of transcript per million mapped reads (FPKM) method was used to calculate the expression levels. Differential expression analysis between two groups (≥ three biological replicates per condition) was performed using the DESeq2 R package (1.14.1). The resulting P-values were adjusted using the Benjamini and Hochberg’s approach for controlling the False Discovery Rate (FDR). Genes with an adjusted P-value <0.05 found by DESeq2 were assigned as differentially expressed.

Gene Ontology (GO) enrichment analysis of differentially expressed genes was implemented by the clusterProfiler R package, in which gene length bias was corrected [50, 51]. GO terms with corrected P-value less than 0.05 were considered significantly enriched by differential expressed genes. KEGG is a database resource for understanding high-level functions and utilities of the biological system, such as the cell, the organism, and the ecosystem, from molecular-level information, especially large-scale molecular datasets generated by genome sequencing and other high-throughput experimental technologies (http://www.genome.jp/kegg/) [52]. We used clusterProfiler R package to test the statistical enrichment of differential expression genes in KEGG pathways.

### 2.9. Real-time qPCR analysis

For standard experiments, groups of approximately twenty embryos were collected and transferred to RNAlater (Qiagen). The following day the RNA was extracted with RNAeasy columns (Qiagen) following the manufacturer’s instructions. For the calcitriol induction experiments, 5-7 halves of embryos were pooled after genotyping. 500 ng was used for reverse transcription with Revertaid M-MLV (Invitrogen) with random primer decamers. cDNA (or DNA for the mitochondrial quantification assay) was diluted 1:40 in 15 µl quadruplicate reactions. The Maxima SYBR Green Reagent (Thermo) was used for qPCR in a Light Cycler 480 (Roche), using the following program: 95 °C, 5 min; (95 °C, 15 s; 58 °C, 20 s; 72 °C, 30 s-single quantifications at this step-) ×50 cycles; and a melting curve from 72 to 95 °C holding during 5 s each 0.5 °C was performed. Relative quantification with the Light Cycler 480 software was performed with at least three of the four replicates that displayed similar reaction curves, after normalizing to the expression level of the elongation factor 1 alpha *(ef1alpha*) and using a double delta Ct method. The primer sequences are listed in supplementary table 1.

### 2.10. Mitochondrial quantification

Mitochondria were quantified on extracted erythrocytes by measuring the number of copies of mitochondrial DNA per genomic DNA. The erythrocytes were extracted as previously described and lysed for DNA extraction. The samples were used for genotyping and diluted 1:100 prior to qPCR quantification. For the absolute quantification of mitochondrial/nuclear DNA ratio, relative values for the mitochondrial gene (*mt-nd1)* & the genomic gene (*pol1*) were compared within each sample to generate a ratio representing the relative level of mitochondrial DNA per nuclear genome. The primers are shown in supplementary table 1.

### 2.11. mRNA synthesis and rescue experiments

The PCMVSPORT3.1 plasmid containing the cDNA of *zmiz1a* (Dr.77379 I.M.A.G.E consortium) was used for mRNA synthesis with the mMessage mMachine SP6 kit (Ambion). Previous to the rescue assays 100, 400, and 800 pg of *zmiz1a* mRNA were separately injected into wild-type embryos to generate a dosage-response curve. The optimal amount of mRNA that did not produce a significant number of dead or defective embryos in comparison to buffer-injected controls was 400 pg. For the subsequent experiments, one-cell stage embryos obtained from heterozygous crosses were injected with 400 pg of *zmiz1a* mRNA.

### 2.12. Vitamin D response assay

24 hpf embryos from a *zmi1a+/-cross* were manually dechorionated and placed in a 24 well plate. Two embryos were placed per well for a total of 24 per condition in either control or 10 μM calcitriol (1,25(OH)2D) solution (Caiman chemical) and incubated for 24 hours. Embryos were collected and cut in half using the tail for genotyping, then pooled according to genotype, and used for RNA extraction and reverse transcription. After cDNA synthesis, the expression levels of *cyp24a1* were determined by qPCR.

## 3. Results

### 3.1. zmiz1 gene in zebrafish

Human ZMIZ1 is homologous to the ZMIZ2 protein whose functions we reported previously for its ortholog during zebrafish development. ZMIZ2 conserves the functional domains described for ZMIZ1, and in the vicinity of the SP-RING/Miz domain, it shares a similarity beyond the Zn finger [53, 54]. This extended domain is known as the X-SPRING (eXtended SP-RING). X-SPRING defines the ZMIZ proteins that are conserved in evolution in several animals from insects to humans. In zebrafish, two *zmiz1* genes encode the Zmiz1a and Zmiz1b proteins. Zmiz1a shares a sequence identity of 83% with human ZMIZ1, whereas Zmiz1b is 64% homologous. The homology between the two Zmiz1 fish proteins is 63%. Because Zmiz1a is more conserved and its expression in embryos is higher, we focused on the functional characterization of this gene. We used a full-length *zmiz1a* cDNA construct to examine the spatiotemporal expression pattern during zebrafish embryogenesis using whole-mount in situ hybridization (WISH) (Supplementary Fig. 1). The earliest detection of *zmiz1a* transcripts was in the neuroectoderm at 75% epiboly. At somite-stages, *zmiz1a* was mainly expressed in the neural regions, tip of the tail and optic primordium. At 24 hpf, *zmiz*1a transcripts were detected in the midbrain, spinal cord, and optic stalk. In addition to these locations, at 36 hpf, we detected additional expression of *zmiz1a* in the floor plate and the pectoral fin buds. Histological sections of embryos at 24, 48 and 72 hpf revealed the presence of *zmiz1a* transcripts in the retina, midbrain and hindbrain, gut and circulatory system (Supplementary Fig. 1H-J).

### 3.2. Loss of zmiz1a led to embryonic death at 15 dpf

To introduce CRISPR/Cas9-mediated mutations in the *zmiz1a* gene, we designed a guide RNA (gRNA) that targeted exon 3 of the reported genomic sequence. Frameshift mutations in this exon created a premature stop codon leading to a truncated protein (80 aa) lacking all functional domains of Zmiz1a except for a small part of the TPR (Fig. 1A). We obtained two mutant alleles with 7- and 22-nucleotide deletions. Crosses of the F1 expanded lines from these alleles produced offspring in which embryos homozygous for the mutation were present at Mendelian frequencies. They did not exhibit any morphological abnormalities. Two groups of *zmiz1a*^*+/+*^ and *zmiz1a*^*-/-*^ fish were compared to test for differences in survival during development. Up to 14 dpf, there was no difference in survival between mutants and wild types. However, at 15 dpf, *zmiz1a*^*-/-*^ larvae showed a dramatic drop in the survival rate compared to *zmiz1a*^*+/+*^ larvae (Fig. 1B). A steady decline in *zmiz1a*^*-/-*^ larvae was detected up to 19 dpf, when 30% of mutant larvae were still alive. Surviving larvae reached adulthood and seemed normal, except for a reduced fertility observed in homozygous females. This survival behavior was the same for both alleles as well as for a heteroallelic combination. Thus, we continued the work with the 22-base pair deletion.

**Figure 1.**
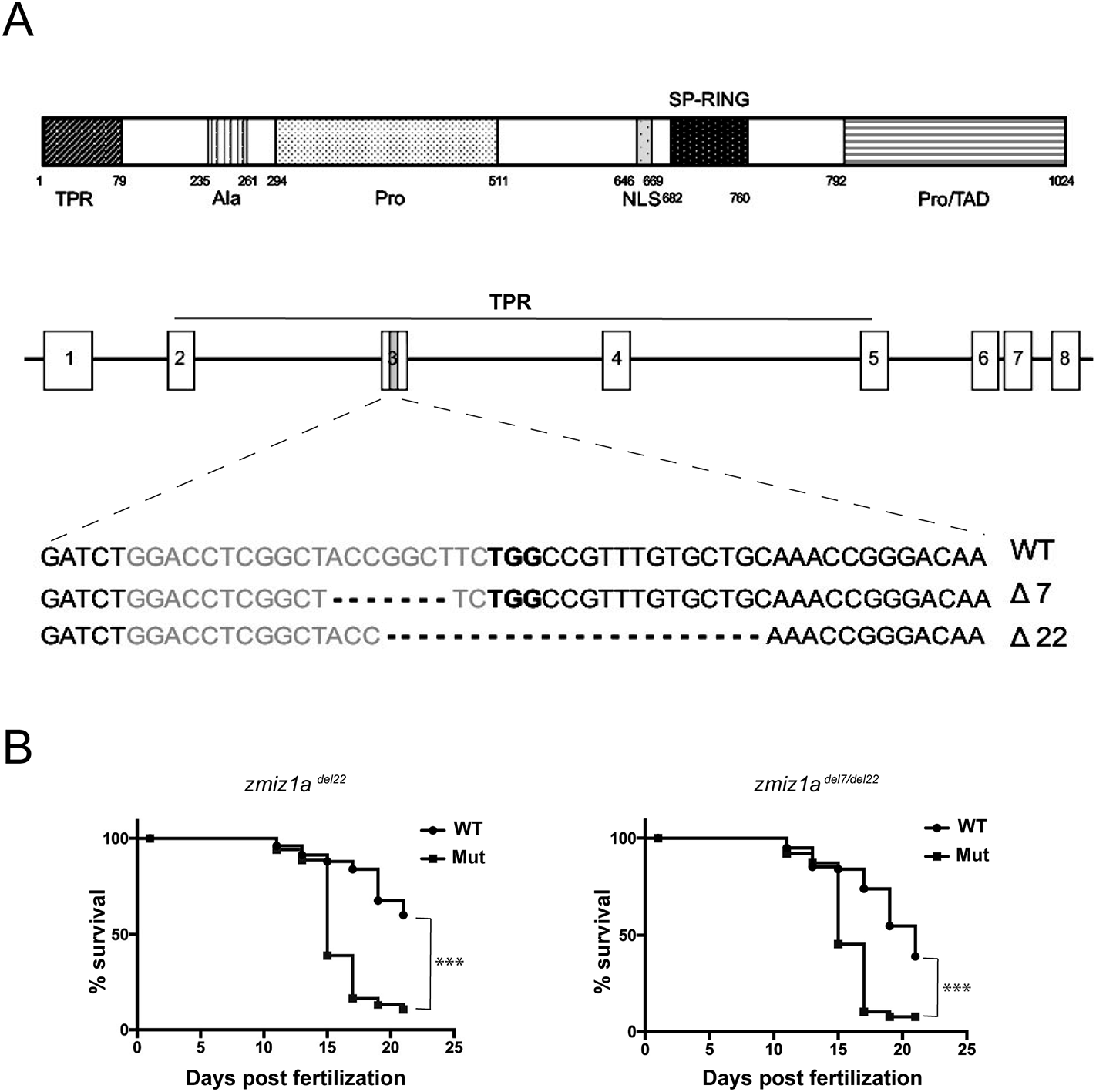
A. Schematic diagram of *zmiz1a* gene. CRISPR/Cas9 mediated-deletions were produced in exon 3. The nucleotide sequence of the RNA-guide targeted region is shown in grey with the PAM in bold and the deletions are indicated in lines below. B. Survival assays for the *zmiz1a* mutant alleles. Percentage survival during the first 25 days of development is shown. Offspring from the heterozygous *zmiz1a*^*del22*^ allele and the *zmiz1a*^*del7/del22*^ allele were compared with that of wild-type progenies. Survival curves are based on data pooled from six individual experiments (*zmiz1a*^*del22*^) and three individual experiments (*zmiz1a*^*del7/del22*^) (*n* > 100 per group in total). The asterisks indicate the significant difference between wild-type and mutant survival (P<0.0001), tested with a log rank test.

### 3.3. zmiz1a mutants have delayed erythrocyte maturation

Since we found reports describing phenotypes of death at similar ages for zebrafish mutants affected by hematopoiesis defects, we analyzed the possibility of an abnormal hematopoiesis in the *zmiz1a*^*-/-*^ fish. Prior to the search for a phenotype in developing hematopoietic cells, we confirmed the presence of *zmiz1a* transcripts in the circulating erythrocytes in embryo sections of WISH experiments at 24 and 48 hpf, and we detected a positive signal in red blood cells (Fig. 2). We then examined erythrocyte development in *zmiz1a* mutants. For this purpose, we collected erythrocytes from larvae after transecting their tails. We compared stage-specific erythrocytes from wild-type and *zmiz1a*-mutants at 3, 7, 11 and 15 dpf. Wright-Giemsa staining of blood smears revealed delayed maturation in the *zmiz1a*-null erythrocytes from 11 and 15 dpf larvae (Fig. 3A). Mutant larvae at these stages presented a percentage of basophilic blood cells with large nuclei resembling proerythroblasts. To quantify the delay of maturation, we measured the nucleus-to-cytoplasm (N:C) ratio and found that this value was significantly higher in the 11 and 15-dpf *zmiz1*-mutated erythrocytes (Fig. 3B).

**Figure 2.**
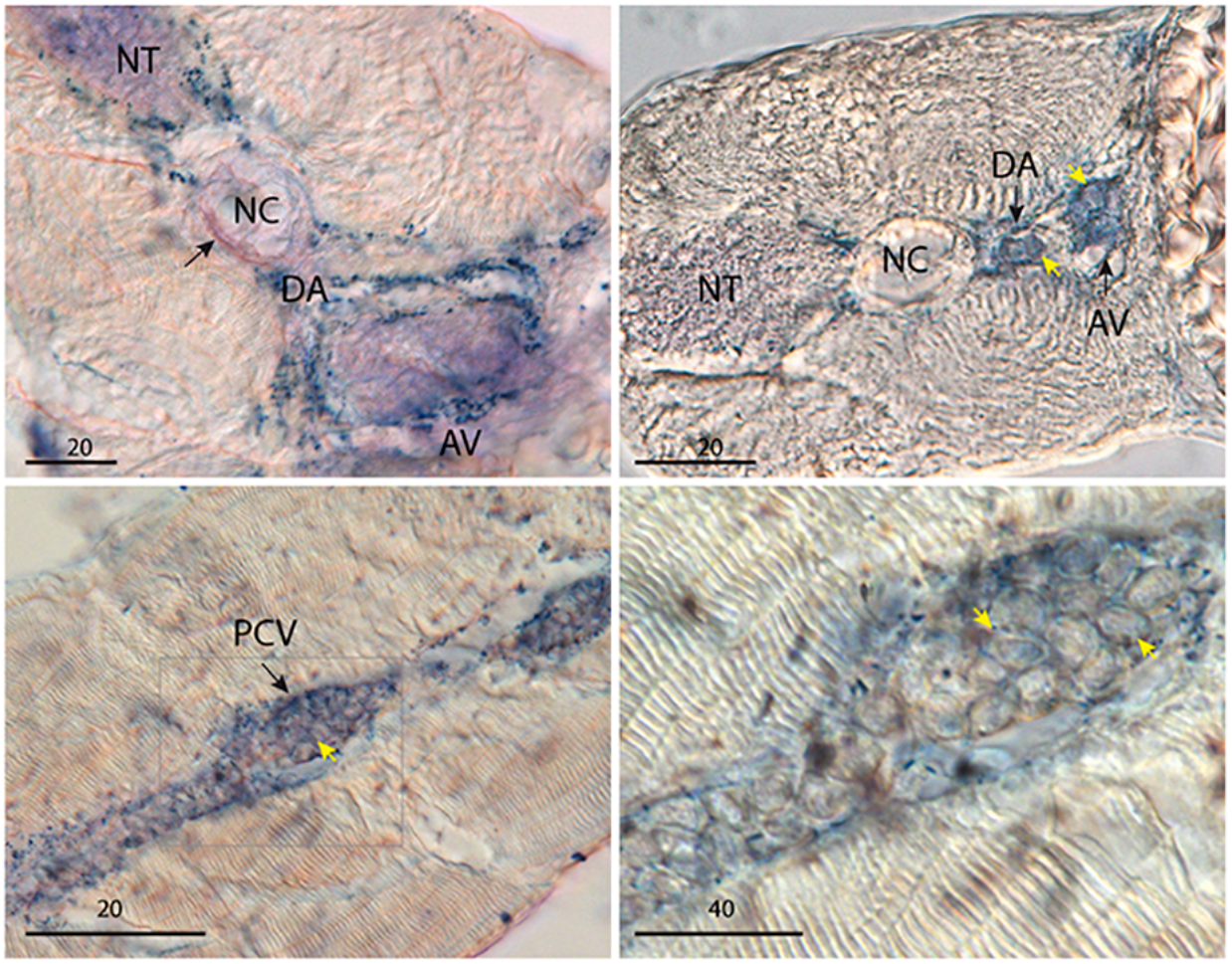
Embryos showing expression of *zmiz1a* in the circulatory system. Sections of embryos after whole-mount in situ hybridization with the *zmiz1a* probe. (upper panels) Sagittal sections of 24 hpf. Scale bars: 20 μm. (lower panels) Sagittal sections of a 48 hpf embryo. (Left) Scale bar: 20 μm. (Right) Scale bar: 40 μm. Yellow arrows indicate a positive *zmiz1a* signal in erythroid cells. AV: Axial Vein, DA: dorsal aorta, NC: notochord, NT: neural tube, PCV: posterior cardinal vein.

**Figure 3.**
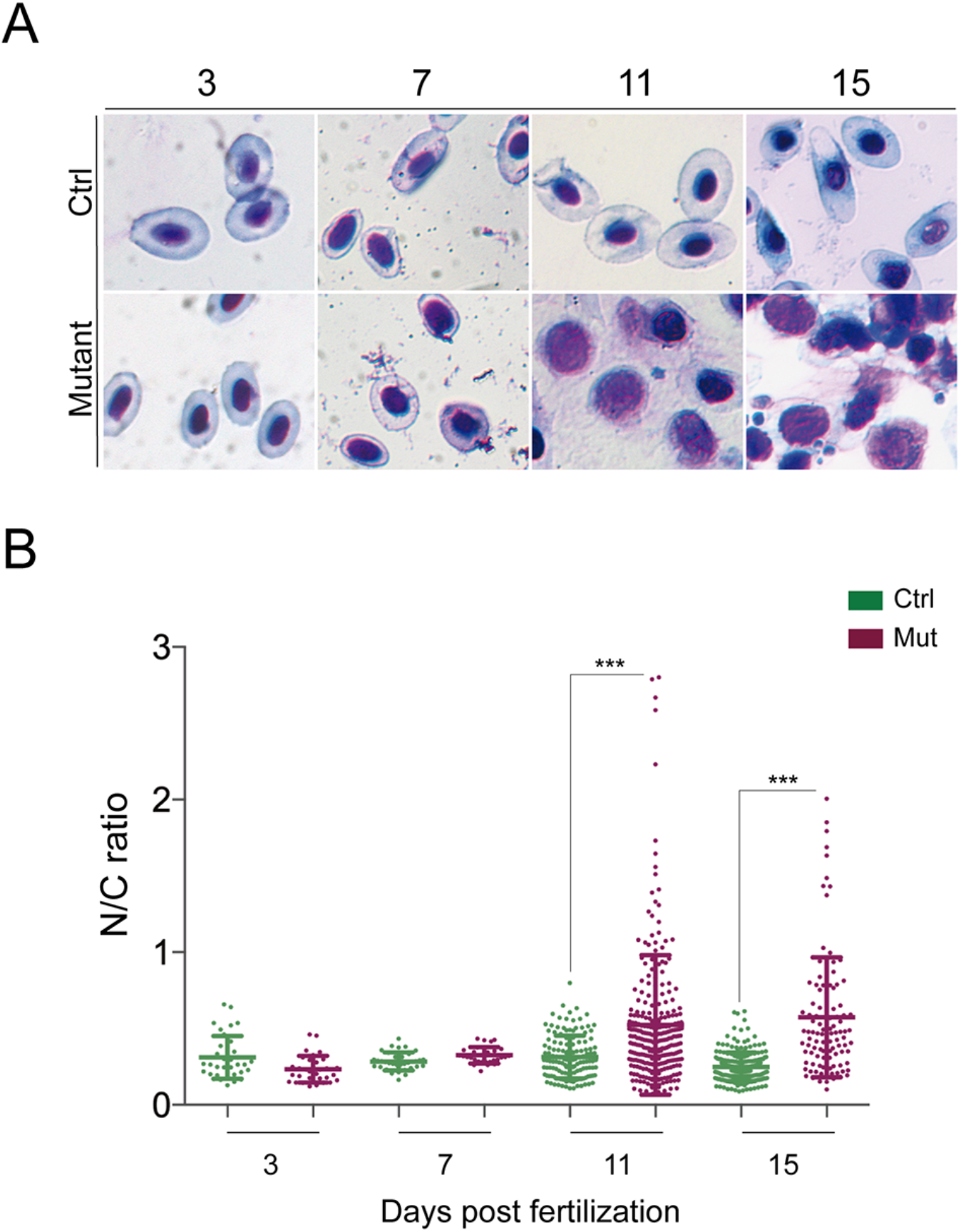
Analysis of circulating erythrocytes from control and *zmiz1a* mutant embryos. A. Wright-Giemsa stained isolated erythrocytes from 3, 7, 11, and 15 dpf larvae. The presence of immature erythroblasts can be appreciated at 11 and 15 dpf. B. Scatter plots representing the tabulation of the N:C area ratio of erythrocytes isolated from control or *zmiz1a*^*-/-*^ 3 to 15 dpf embryos. Horizontal lines indicate mean values. Data are mean ± 1SD for > 110 cells. P < 0.0001.

To evaluate whether the zmiz1-nule embryos carried hemoglobin deficiencies, we stained isolated erythrocytes from wild-type and mutant 15 dpf embryos with *O*-dianisidine (*O*-das) and found that they all showed similar intensities, which indicated a normal hemoglobinization (Fig. 4A). However, whole embryo *O*-das staining (at 48 hpf) showed that while the intensity of the signal was similar in wild-type and mutant embryos, there was a difference when we measured the yolk-stained area, which was significantly smaller in the *zmiz1a*-mutant embryos (Fig. 4B). A possible interpretation of this result is that circulating erythrocytes are distributed differently along the embryo or in the yolk in the control and mutant embryos.

**Figure 4.**
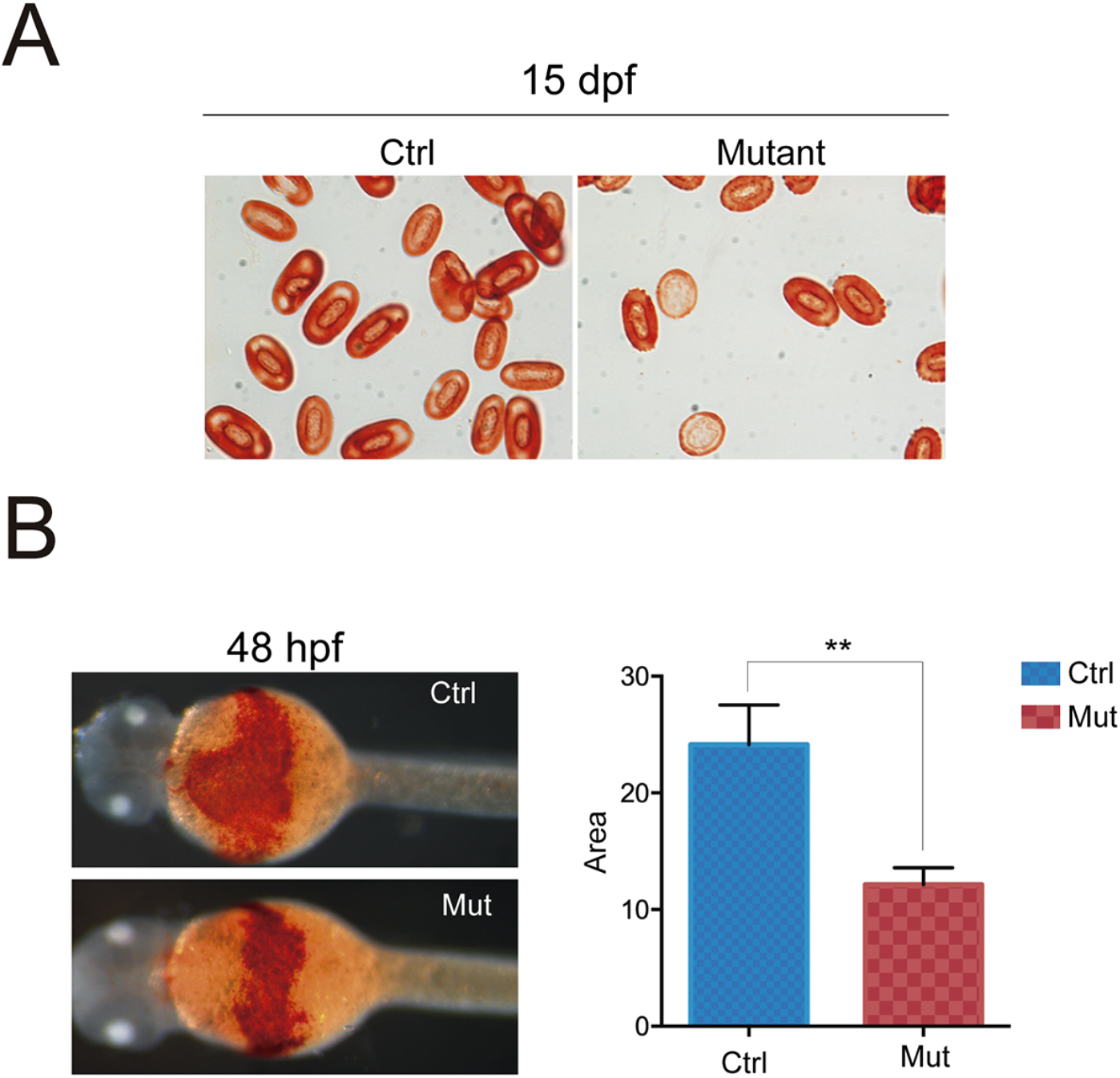
Analysis of hemoglobinization of erythrocytes in the *zmiz1a* mutant embryos. A. O-dianisidine (O-das) staining of isolated blood cells from 15 dpf control and *zmiz1a*^*-/-*^ embryos shows a normal hemoglobinization of mature erythrocytes in the mutants and a weaker signal in immature erythroblasts. B. O-das staining of 48 hpf whole embryos showing a reduced stained area in the yolk of the *zmiz1a*^*-/-*^ embryos. Data are mean ± 1SD for ≥13 embryos. P < 0.001

### 3.4. Primitive hematopoiesis and HSC specification are normal in zmiz1a mutants

To analyze whether primitive hematopoiesis was defective in the *zmiz1a* mutant embryos, we examined the expression pattern of *gata1* in embryos at 18 to 30 hpf (Supplementary Fig. 2); we found that *gata1* is normally expressed at early stages. At 30 hpf, mutant embryos exhibited substantial numbers of positive cells throughout the yolk sac, which were not detected in control embryos (Supplementary Fig. 2E-F). This increased signal could reveal the presence of migrating *gata1*-expressing progenitors in the mutants and it might be correlated with the abnormal *O*-das staining found in 48 hpf mutant embryos. Apart from this observation, the specification of primitive erythrocytes seems normal in the *zmiz1a* mutants. To analyze if hematopoietic stem cells were produced in the *zmiz1a* mutant embryos, we examined *runx1* expression in 30 hpf and 5 dpf embryos. WISH revealed a normal *runx1* pattern in the *zmiz1a* mutants, indicating the presence of HSCs (Supplementary Fig. 2G-J). To further test whether definitive hematopoietic cells in the *zmiz1a* mutants were able to undergo lymphoid differentiation, we examined *rag1* expression at 5dpf. Both, control and mutant embryos showed a *rag1* positive signal in the thymus, indicating that a proper T-cell development occurred in the *zmiz1* embryos (Supplementary Fig. 3A-B). Finally, we assessed the formation of myeloid cells in *zmiz1*^*-/-*^ embryos. For this purpose, we used zebrafish transgenic lines with green fluorescent neutrophils expressing *Tg(mpx:eGFP)*^*i114*^ or red fluorescent macrophages expressing *Tg(mpeg1:mCherry)*^*gl22*^ to visualize these cells in control and mutant embryos. We found an apparently normal abundance of both neutrophils and macrophages in 4pdf *zmiz1a* embryos compared with control embryos (Supplementary Fig. 3C-F). Altogether, our analysis showed that the *zmiz1a* mutants have a normal primitive and definitive hematopoiesis and proper myeloid and lymphoid differentiation, which indicates that the defects observed during the maturation of erythrocytes are independent of these processes.

### 3.5. RNA-seq analyses revealed the downregulation of autophagy genes in the Zmiz1a mutant

To develop a comprehensive characterization of Zmiz1-mutant embryos, we examined global differences in gene expression. To this aim we conducted transcriptome sequencing (RNA-seq) analysis of 15 dpf *zmiz1a*-mutant and wild-type embryos. Pearson correlation between samples (Supplementary Fig. 4A) and heat map analysis (Fig. 5A) showed clustering of three mutant and four wild-type biological replicates of each population, demonstrating that the two embryo groups are distinct. Differential expression analysis uncovered 2892 differentially expressed genes (DEGs), of which 1309 were upregulated and 1583 were downregulated (Fig. 5B). Through Gene Ontology (GO) classification, we found that DEGs were enriched for the biosynthesis and metabolism of sterols and steroids, mitochondrial organization and autophagy (Fig. 5C). In the cellular component terms, the *zmiz1a* mutation was significantly associated with the endoplasmic reticulum, mitochondria and autophagosomal components (Supplementary Fig. 4B). Notably, the autophagy genes were mostly downregulated (Fig. 5E). This is evident in the Kyoto Encyclopedia of Genes and Genomes (KEGG) analysis, where we found that the top enriched pathways for the downregulated DEGs were those related to animal autophagy (Fig. 5D).

**Figure 5.**
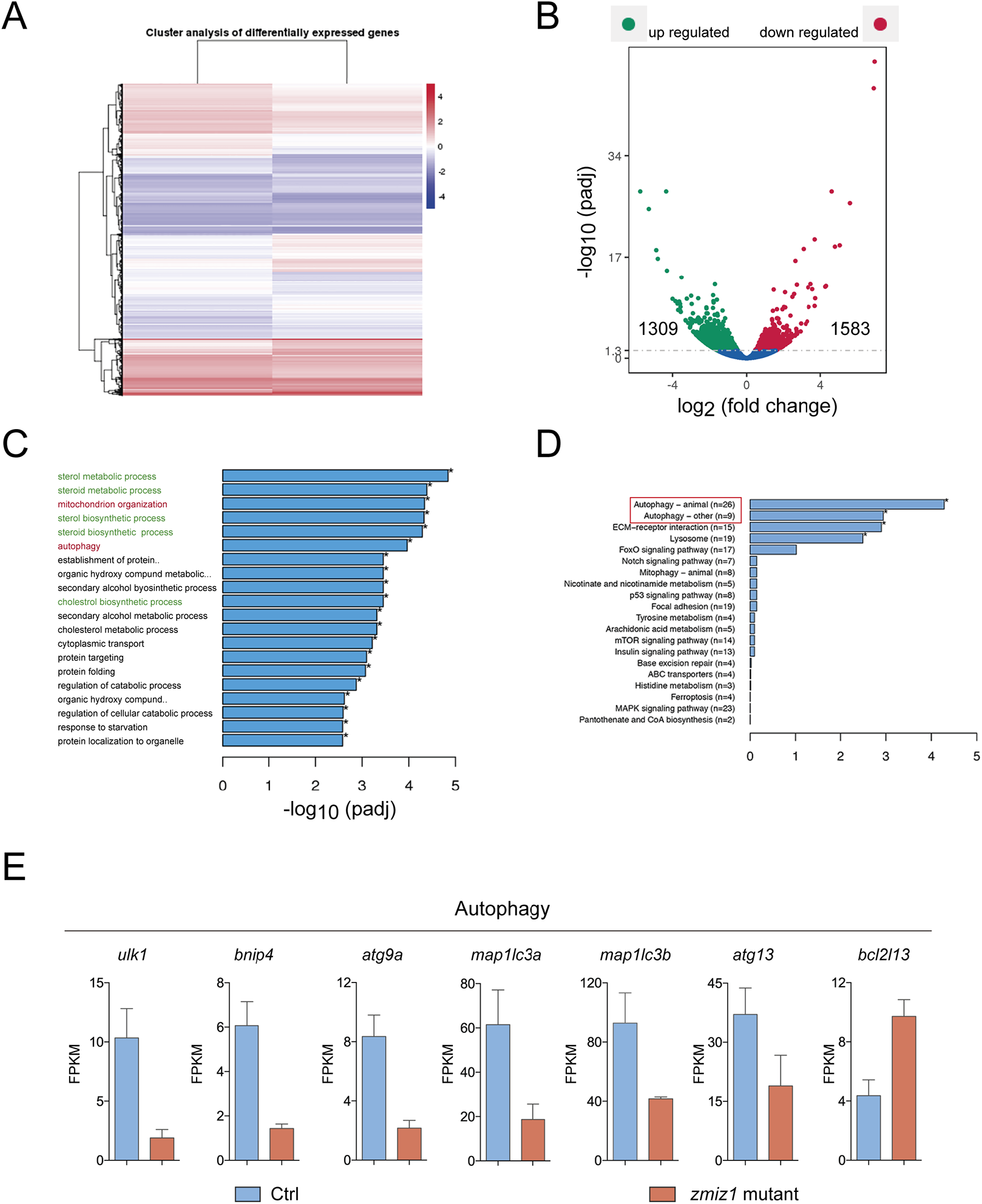
Transcriptome analysis of 15 dpf control and *zmiz1a*^*-/-*^ whole embryos. A. Hierarchical Clustering Heatmap of gene expression data determined for four replicate control and three replicate mutant samples. The overall results of FPKM (Fragments Per Kilobase of transcript sequence per Millions of base pairs sequenced) cluster analysis, clustered using the log10 (FPKM+1) value. Red denotes genes with high expression levels, and blue denotes genes with low expression levels. The color ranging from red to blue indicates log10 (FPKM+1) value from large-to-small. B. Volcano plot. Horizontal axis for the fold change of genes in the control and mutant samples. Vertical axis for statistically significant degree of changes in gene expression levels, the smaller the corrected pvalue, the bigger -log10 (corrected pvalue), the more significant the difference. The point represents one gene, blue dots indicate no significant difference in genes, red dots indicate upregulated differential expression genes, green dots indicate downregulated differential expression genes. C. GO enrichment bar plot for Biological Process (BP). The top 20 significantly enriched terms in the GO enrichment analysis. D. KEGG enrichment bar plot. The top 20 enriched terms in the KEGG enrichment analysis showing four with significant corrected pvalues. E. Bar charts comparing the normalized average expression levels (FPKM) for autophagy genes *ulk1a, atg9a, bnip4* and *atg13* in control and *zmiz1a* mutant samples.

The RNA-seq data involving autophagy and erythropoiesis genes were validated by real-time polymerase chain reaction (qPCR). This analysis confirmed the downregulation of *ulk1a, bnip4, atg9a* and *atg13* in the Zmiz1a-mutants compared to control larvae (Fig. 6A). Autophagy is a catabolic pathway that is the main mechanism responsible for the removal of organelles during red blood cell development. In particular, mitochondrial clearance by mitophagy is an essential step for terminal erythroid differentiation [55, 56]. Mice with deletions in autophagy genes retain mitochondria in mature erythrocytes and display anemia and reticulocytosis [57]. In zebrafish mitophagy is also the primary method used to eliminate mitochondria from erythroblasts [58]. To evaluate the clearance of mitochondria in the *zmiz1a* mutants, we determined the ratio between mitochondria (mt-DNA) and nuclei (n-DNA) in circulating erythroid cells isolated from 15 dpf larvae. For this purpose, we used qPCR to measure the levels of the NADH dehydrogenase 1 (*nd1)* gene, encoded in the mitochondrial genome, which we normalized to the DNA levels of the nuclear-encoded DNA polymerase gamma (*polg1)* gene [59]. This analysis indicated an increased mt-DNA copy number in the *zmiz1a* mutants relative to the wild-type blood cells (Fig. 6B). This result, together with the reduced expression of autophagy genes in the mutant embryos, suggests that the delayed maturation detected in the red blood cells (RBCs) of the *zmiz1a* mutant could be due to impaired mitophagy which leads to the retention of mitochondria.

**Figure 6.**
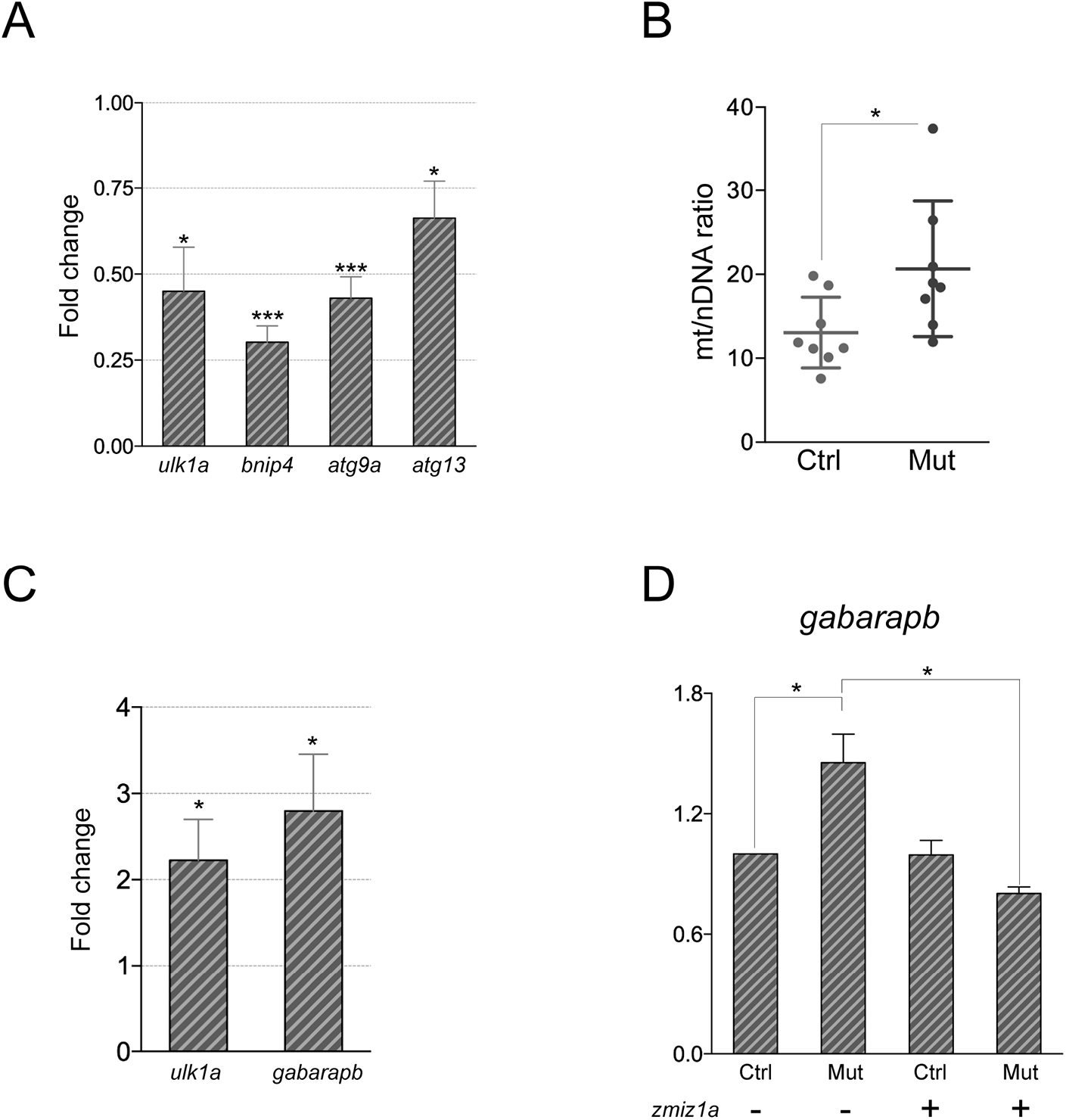
Analysis of autophagy gene expression and mitochondrial DNA copy number. A. Quantification by qPCR of autophagy gene expression. 15 dpf embryos were evaluated. For all genes n ≥ 3. *P< 0.05, ***P<0.001. B. Quantification of mitochondrial DNA copy number in isolated erythrocytes from control and *zmiz1a* mutant 15 dpf larvae. Each point represents the mitochondrial/nuclear DNA ratio determined for the erythrocyte population isolated from one embryo. Mitochondrial and genomic DNA is used for qPCR quantification of *nd1* and *polg1* respectively. Horizontal lines indicate mean values. Data are mean ± 1SD for 8 embryos. P < 0.05. C. Quantification by qPCR of the expression levels of autophagy genes *ulk1a*, and *gabarapb* in 48 hpf embryos. For both genes n ≥ 3. P< 0.05. D. Graph showing a rescue experiment with injected *zmiz1a* mRNA. Quantification of *gabarapb* gene expression shows that the upregulation of *gabarapb* in the mutant embryos is compensated by the injection of the *zmiz1a* mRNA. n= 4. *P< 0.05.

To determine whether the altered expression of autophagy genes represented a general defect in the *zmiz1* mutants not specifically related to the removal of organelles during RBCs maturation, we analyzed the expression of autophagy genes in 48 hpf mutant and wild-type embryos. We found significantly altered expression of autophagy genes, although with the opposite tendency, with the upregulation of *ulk1a* and *gabarapb* (Fig. 6C). To demonstrate the specificity of the *zmiz1a* knockout phenotype, we performed a rescue experiment to prove that the alterations in gene transcription found in the mutant embryos were due to the loss of Zmiz1a function. We injected single-cell embryos obtained from *zmiz1a*^+/-^/ *zmiz1a*^-/-^ crosses with synthetic *zmiz1a* mRNA and collected them at 48 hpf. After genotyping, pools of *zmiz1a*^+/-^ and *zmiz1a*^-/-^ embryos were used for quantification of *gabarapb* gene expression. We found that *zmiz1a*^-/-^ embryos injected with *zmiz1a* RNA recovered *gabarapb* expression levels, which were upregulated in the mutants (Fig. 6D). Thus, *gabarapb* upregulation is a consequence of a lack of Zmiz1a activity. Collectively our results show that Zmiz1a is required for the correct expression of the autophagy genes in zebrafish.

### 3.6. The zmiz1a mutants exhibit alterations in proinflammatory gene expression and a reduced response to vitamin D

Autophagy is known to contribute to the regulation of inflammation and immune system function. In particular, impaired mitophagy causes mitochondrial dysfunction, increased oxidative stress and susceptibility to proinflammatory stimuli [60, 61]. In light of this information, we decided to quantify proinflammatory gene expression in the *zmiz1a* mutant fish. We performed qPCR to compare the expression of the inflammatory cytokines *il1b, Il6, caspase-1* and *tnfa* in 48 hpf mutant and control embryos. We found a significant upregulation of the *il1b* gene in the *zmiz1a*^*-/-*^ embryos, while *tnfa* exhibited a significant downregulation (Fig. 7A). This result suggests that the *zmiz1a* mutant fish might have abnormal inflammatory responses.

**Figure 7.**
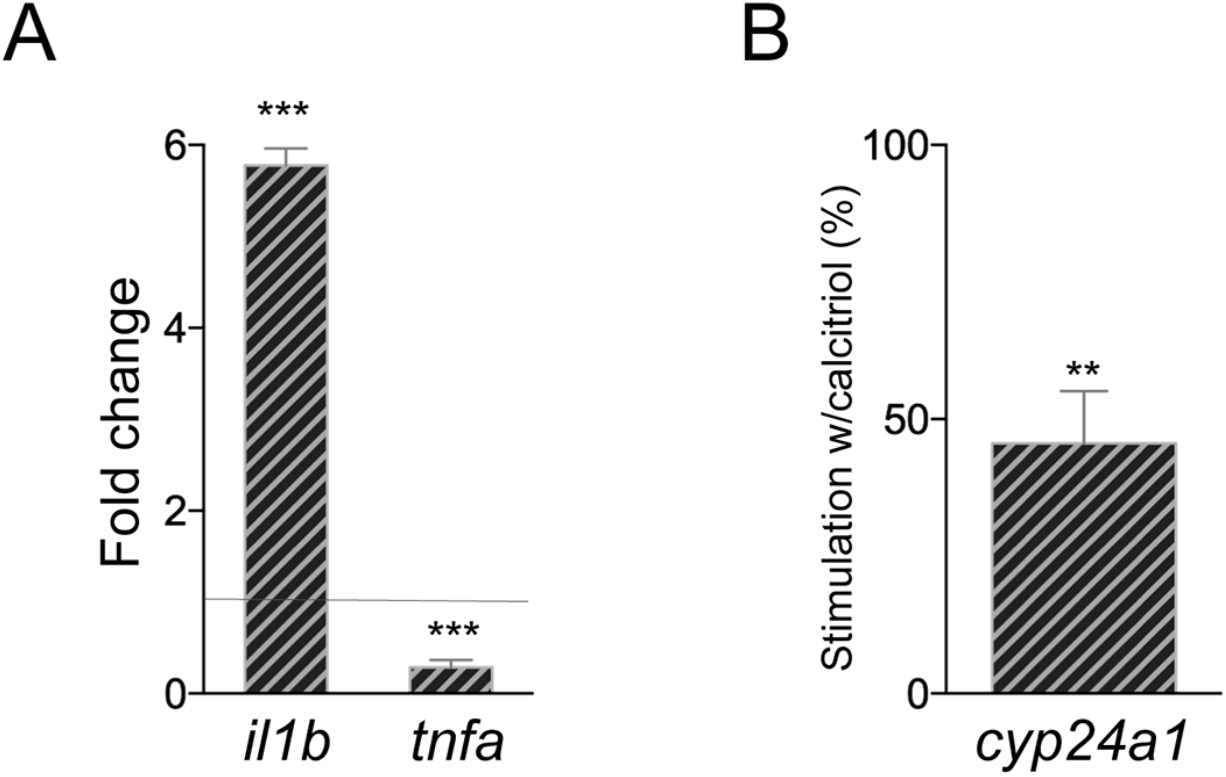
Proinflammatory gene expression and response to vitamin D. A. Quantification by qPCR of the expression levels of proinflammatory genes *il1b* and *tnfa* in 48 hpf embryos. For both genes n ≥ 3. ***P<0.001. B. Effect of exogenous calcitriol on *cyp24a1* induction in 48 hpf embryos. The graph shows the percentage of stimulation of *cyp24a1* transcription relative to 100 percent assigned to the control embryos, after incubation with calcitriol. Value is the mean ± 1SD; n=4. P< 0.01.

Human ZMIZ1 is a LDAD risk gene. In particular for multiple sclerosis (MS), both ZMIZ1 gene transcription and protein levels are reduced in blood in people with MS, and ZMIZ1 is regulated by vitamin D in monocytes [22]. The vitamin D receptor (VDR) binds to superenhancer regions in the *ZMIZ1* gene in immune cells, and single nucleotide polymorphisms (SNPs) associated with risk were mapped to the VDR binding peaks [21]. These results support the use of ZMIZ1 gene expression as a biomarker for MS. Furthermore, it was suggested that higher expression of ZMIZ1 in response to vitamin D results in protection across LDADs. Therefore, the manipulation of ZMIZ1 expression in a model system could be a useful tool for the study of LDADs pathogenesis.

In our RNAseq study, the top terms of the GO BP analysis included the enrichment of “steroid biosynthesis”, and some DEGs in this term were related to the vitamin D pathway, including *pth1a, tmsf2, ebp, hsd17b7, sc5d*, and *fbp1b* (Supplementary Fig. 4C). Considering these data, we decided to evaluate the response to vitamin D in live zebrafish *zmiz1a* mutants. To visualize vitamin D signaling activity, we incubated 24 hpf control and mutant embryos in the presence of 1,25(OH)2D3 (calcitriol), which is the active hormone derived from vitamin D that binds to the VDR. After one day of treatment, we quantified the expression of *cyp24a1*. Cyp24a1 is the cytochrome 450 enzyme that specializes in the degradation of calcitriol and is the most highly induced vitamin D target gene; thus, it is a good indicator of the vitamin D signaling activity [62]. We found that in the control embryos, calcitriol induced up to a hundred-fold change in the expression of *cyp24a1*. This stimulation was greatly reduced in the *zmiz1a* mutants, which showed 60% less stimulation of *cyp24a1* expression compared to control embryos (Fig. 7B). This result indicates that Zmiz1a is required for an optimal response to vitamin D.

## 4. Discussion

In this study, we analyzed the phenotype produced by the loss-of-function of *zmiz1a* gene in zebrafish. ZMIZ1 is a transcriptional coactivator that contains a variety of structural domains. These features enable a diversity of molecular interactions that could allow this protein to participate in different biological functions. We found that *zmiz1a* mutant embryos at 11 dpf had immature erythroblasts in their blood, which indicates a defective terminal differentiation of their definitive erythrocytes. Our RNA-seq study uncovered the downregulation of the autophagy genes *ulk1, atg13, bnip4*, and *atg9a*. Ulk1 is a serine-threonine kinase that lies on top of the autophagy pathway and initiates both canonical and ATG-5-independent autophagy. In mammals, Ulk1 plays a critical role in the clearance of mitochondria and ribosomes during the differentiation of erythroblasts, and *ulk1* expression levels correlate directly with the removal of mitochondria by autophagy. Therefore, the decreased expression level of *ulk1* in the *zmiz1a* mutant fish could be a cause of the delayed maturation of the erythroblasts. Studies based on the use of human erythroblast cultures show that autophagy is induced at the polychromatic erythroid stage and is accompanied by the transcriptional upregulation of several autophagy genes. GATA1 was identified as a critical factor that directly activates the transcription of autophagy genes [63]. Human ZMIZ1 is included among the GATA1 target genes in *the autophagy regulatory network* (ARN) database. A plausible hypothesis is that Zmiz1a interacts with Gata1 and acts as a coactivator during the upregulation of autophagy genes. Also, p53 is well known for its role as a transcription factor that regulates autophagy. In fact, several genes encoding proteins involved in autophagy are known to be direct p53 targets. Thus, Zmiz1a could also participate in the regulation of autophagy through its function as a coactivator of p53. In our search for abnormal gene expression in 48 hpf Zmiz1 mutant embryos, we observed dysregulation of the autophagy genes *ulk1a* and *gabarapb*. Interestingly, at this stage, these genes were upregulated. Altogether, the consistent alteration of the expression of autophagy genes at two different stages of development suggests that an abnormal autophagy response might be associated with *zmiz1a* loss-of-function and that Zmiz1a is involved in the regulation of Ulk1 or other autophagy genes.

A percentage of Zmiz1a mutant fish survive to adulthood and seem normal. This result raises the question of why adult fish might have milder or null effects in their erythropoiesis. In mice, elimination of mitochondria from embryonic reticulocytes involves the alternative Ulk1-dependent Atg5-independent macroautophagy pathway; however, primitive and adult definitive reticulocytes do not depend on this pathway for mitochondrial clearance [64, 65]. In zebrafish, there is yet no evidence indicating the existence of the alternative autophagy pathway, nor is anything known about the types of macroautophagy operating in erythroblasts during specific stages of life. The lack of a visible phenotype in the Zmiz1a mutant fish could reflect that a different mechanism of mitophagy operates at adult stages, but it could also be indicative of the high capacity of erythroid cells to adapt to diminished autophagic activity during terminal differentiation.

Although human ZMIZ1 is known to coactivate transcription mediated by AR, p53, and SMAD3 *in vitro*, only its role in Notch signaling has been well described *in vivo*, in a conditional mouse model in which Zmiz1 was shown to act as a cofactor of Notch1 during thymocyte development and drive pre-T cell proliferation [18]. Zmiz1 cooperates with Notch1 to induce the Notch target genes *Hes1, Lef1*, and *Myc*. In this work, we found that zebrafish *zmiz1a* mutant embryos have normal expression of *rag1* in the thymus, which indicates the presence of early T-lineage progenitors. However, in our study, we did not follow the development of thymocytes during their progression throughout the different CD4-CD8 negative and positive stages in which Zmiz1 is also required for the Notch-dependent double-negative double-positive transition [66]. Thus, the participation of Zmiz1a in zebrafish Notch-dependent thymocyte differentiation cannot be discarded. Zebrafish is a good model to analyze this question given that adaptive immune-deficient fish can reach adulthood, and unlike mouse models, can be maintained under conventional conditions [67]. Related to the coactivation of Notch1 target genes, we did not find alterations in the expression of Notch target genes in the transcriptome of 15 dpf *zmiz1a* mutant fish or in a qPCR analysis that we performed in embryos at 48 hpf. This finding indicates that Zmiz1a is not a general coactivator of Notch signaling. However, these studies were performed using whole embryos, and therefore, if the coactivation effect of Zmiz1a is context-dependent it could be undetected.

We also found alterations in proinflammatory gene expression with the upregulation of *il1b* and the downregulation of *tnfa*. These findings are consistent with an altered inflammatory response in *zmiz1a* mutant embryos. Significantly, the reduced expression of ZMIZ1 in human monocytes has been associated with an increased tendency toward chronic inflammation. Inflammation is also relevant for hematopoiesis and during the terminal differentiation of erythrocytes. Tyrkalska *et al*. reported that in zebrafish, the inflammasome regulates the erythroid-myeloid decision in HSCs and the terminal differentiation of erythrocytes, both events occurring through the cleavage of GATA1 by Caspase-1 [68]. In our study, we did not find differential transcription of the Caspase-1, and we did not detect alterations in the number of neutrophils in 4 dpf mutant embryos; however, it would be interesting to determine the erythrocyte/neutrophil ratio and the activity of Caspase-1 in the *zmiz1a* mutants.

Finally, we discovered that Zmiz1a contributes to the response to vitamin D. This finding is significant because of the extensive literature that indicates a protective role of ZMIZ1 against autoimmunity. Besides, gene clusters whose expression variations correlate with the ZMIZ1 expression pattern in MS patients appear to be regulated by induced vitamin D superenhancers [21]. Interestingly, our result suggests the possibility of reciprocal regulation of ZMIZ1 expression and vitamin D response, implicating that the role of ZMIZ1 in autoimmunity is more important than it was considered. Thus, it will be important to determine if there are physical interactions between VDR and Zmiz1a and if they bind together to a particular set of genes. Several of the most upregulated DEGs identified in our RNAseq study are involved in cholesterol and vitamin D biosynthesis. This overexpression could be the result of a compensatory reaction to a VDR dysfunction. Metabolism and autophagy are both physiological processes that need to be carefully regulated to maintain cellular homeostasis. Imbalance of these processes leads to inflammation. Zmiz1a seems to affect the coordination of all three processes. However, there is still a lot to learn from the *zmiz1a* mutant fish. We do not yet know the molecular actions of Zmiz1a during the regulation of gene expression. Zmiz1a could operate as a coactivator that facilitates the binding of transcription factors such as Gata1, p53 or Vdr, or it could function as an E3-SUMO ligase and mediate the sumoylation of these TFs. Another relevant question is whether Zmiz1a has independent functions in autophagy and the vitamin D response or whether the alterations in these signaling cascades are secondary to Zmiz1a central activity. The *zmiz1a* mutants could be a good zebrafish model for the studying the modulation of the inflammasome, the effects of vitamin D, and its relation with disease and autoimmunity.

In conclusion, in this work, we show that Zmiz1a is required for the maturation of embryonic definitive erythrocytes; we provide evidence of altered autophagy gene expression in 15 dpf larvae, which potentially could cause retention of mitochondria in circulating erythroblasts; and we demonstrate that Zmiz1a functions to stimulate the response to vitamin D in embryos.

## Acknowledgments

We thank Drs. Leonard Zon, Katherine Crosier and Catherine Willett for kindly providing plasmids for ISH; Dr. Herman Spaink for the donation of zebrafish transgenic lines. We thank the IBT/UNAM synthesis facility for oligonucleotide synthesis, Dulce Pacheco for help with fish maintenance, and the librarian Shirley Ainsworth.

## Funding

This work was supported by DGAPA-UNAM grant IN200519 and CONACyT grant 1755. F. Castillo received the CONACyT scholarship 436027.

## Credit author statement

Francisco Castillo: Formal Analysis and Investigation. Laura Ramírez: Investigation. Hilda Lomelí: Conceptualization, Formal Analysis, Visualization, Supervision, Project administration, Funding, Writing-Review and Editing.

## Notes

### Competing Interest Statement

The authors have declared no competing interest.

