## Supplemental material for "*zmiz1a* zebrafish mutants have defective erythropoiesis, altered expression of autophagy genes, and a deficient response to vitamin D"

| gene | Purpose | orientation | sequence |
|---|---|---|---|
| *zmiz1a* | cloning | forward | TTCTCCACAGGTTGTAAGCC |
| *zmiz1a* | cloning | reverse | GACTGCTGGAAAGTGAACTC |
| *zmiz1a* | genotyping | forward | TTCTCCACAGGTTGTAAGCC |
| *zmiz1a* | genotyping | reverse | AAGATTTCCCGTCACATCAC |
| *ulk1* | qpcr | forward | GAAGCTCCAAAGTCCCTCCC |
| *ulk1* | qpcr | reverse | ACTGGCTCACTGGTGAACTG |
| *bnip4* | qpcr | forward | TGCTTGGATTGGGGATCT |
| *bnip4* | qpcr | reverse | GGAGGAAAACAAAAGGCT |
| *atg9a* | qpcr | forward | TGGCCAAGAATGTGGCGTTT |
| *atg9a* | qpcr | reverse | TGTCTGGGATAAACGACCTGC |
| *atg13* | qpcr | forward | AGCCCACAGGACAAGAAAC |
| *atg13* | qpcr | reverse | GGTGATGAAGACGAGCAAG |
| *mt-nd1* | qpcr | forward | CCCACGATTCCGATACGAC |
| *mt-nd1* | qpcr | reverse | GTGCGATTGGTAGGGCGAT |
| *pol1* | qpcr | forward | CTAGGGCTCCAAGATGCGT |
| *pol1* | qpcr | reverse | GGCGTTTCATGTGCTCCTT |
| *gabarapB* | qpcr | forward | CGTCATTCCCCCTACTTCCG |
| *gabarapB* | qpcr | reverse | GGGTGTGTTTTCCCTTTGGC |
| *cyp24a1* | qpcr | forward | GATACCGTGCTGGGCGATTA |
| *cyp24a1* | qpcr | reverse | CCAAACGGCACATGAGCAAA |
| *il1b* | qpcr | forward | GAACAGAATGAAGCACATC |
| *il1b* | qpcr | reverse | ACGGCACTGAATCCACCAC |
| *tnfa* | qpcr | forward | TCCAAGGCTGCCATCCATT |
| *tnfa* | qpcr | reverse | CAAAGACACCTGGCTGTAG |

Supplementary Table 1.
List of primers used for cloning or genotyping the *zmiz1a* mutants, and for quantitative qPCR.

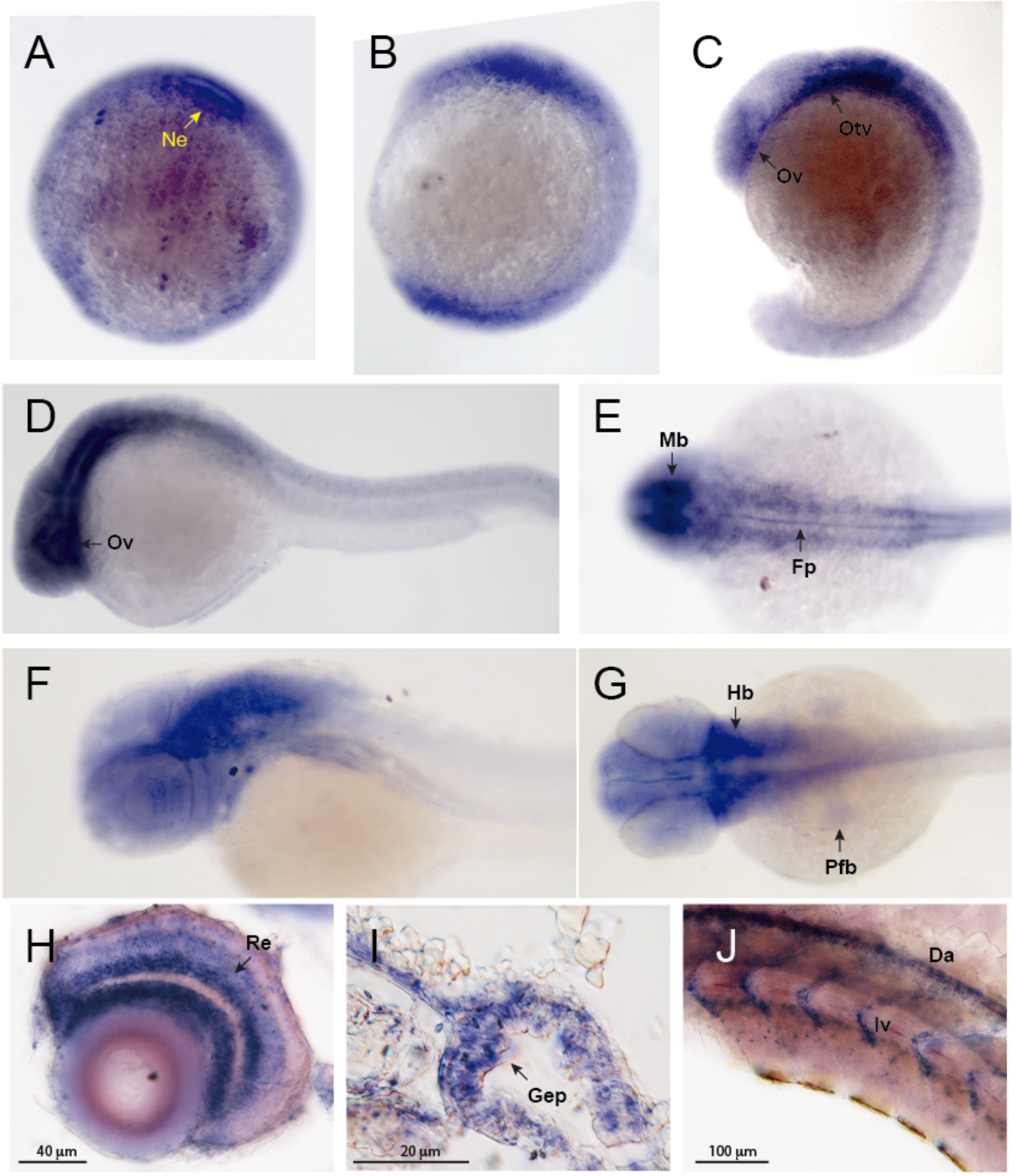


Supplementary Figure 1.

Expression pattern of *zmiz1a* detected by whole-mount in situ hybridization. A. Lateral view of a 75%-epiboly embryo with the animal pole oriented at the top; B-C. Lateral views of embryos at 3- and 18-somite stage respectively, with anterior at the top. D-E. 24 hpf embryos. D. Lateral with anterior oriented towards the left. E. Dorsal view with anterior at the left. F-G. 36 hpf embryos. F. Lateral with anterior toward the left. G. Dorsal with anterior at the left. H-J. Histological sections of embryos. H. Transverse section of 72 hpf retina. I. Transverse section of 72 hpf embryo showing expression in the gut epithelium. F. sagital section of a 48 hpf embryo with *zmiz1a* signal in the vascular system. Da: dorsal aorta, Iv: intersegmental vessel, Fp: floorplate, Gep: gut epithelium, Hb: hindbrain, Mb: midbrain, Ne: neuroectoderm, Ov: optic vesicle, Otv: otic vesicle, Pfb: pectoral fin-bud, Re: retina,

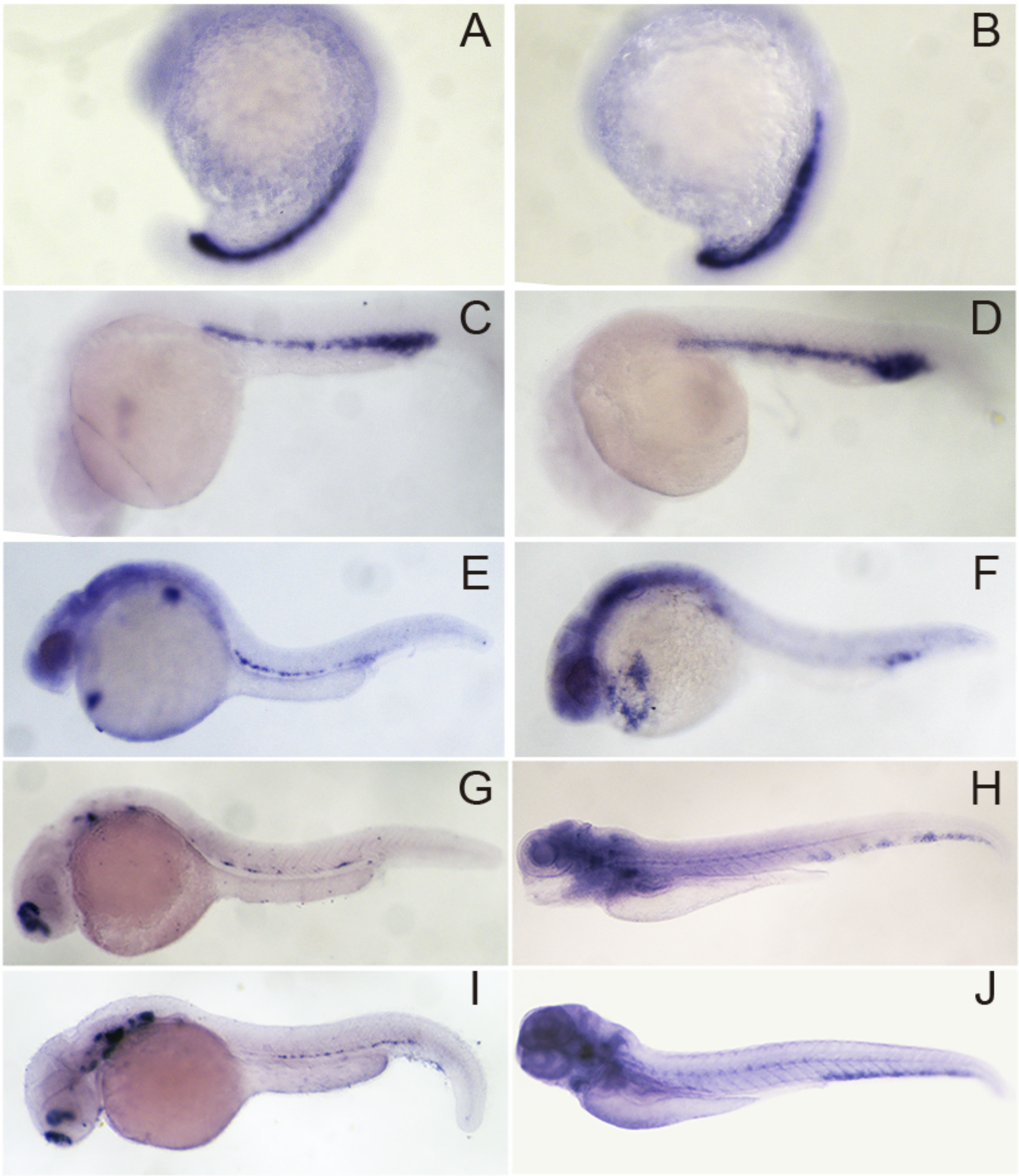


Supplementary Figure 2.

Expression of hematopoietic markers in wild type and *zmiz1a* mutant embryos. Lateral views of embryos showing expression of *gata1* (A-F), and *runx1*(G-J). A, C, E, G and H. Control embryos at the indicated stages. B, D, F, I and J. *zmiz1a* mutant embryos. Arrow in F indicates additional positive signal in the yolk of the mutant embryos.

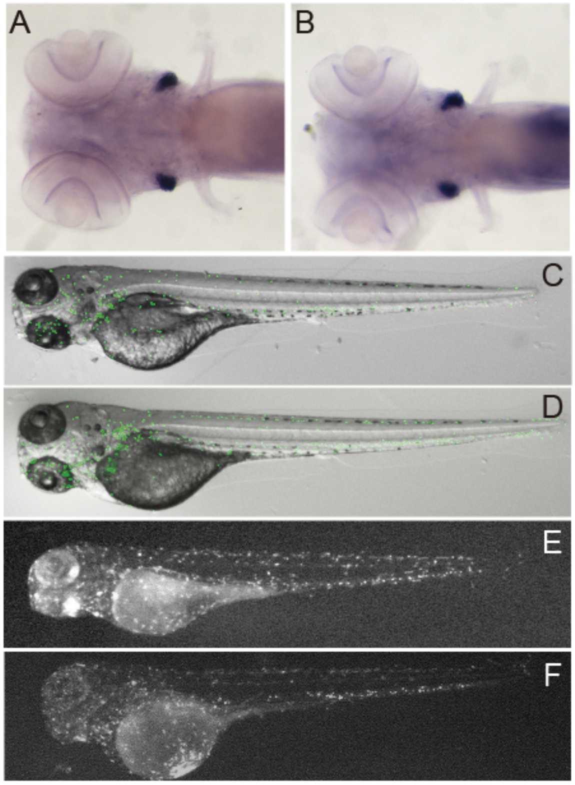


Supplementary Figure 3.

Expression of myeloid and lymphoid markers in wild type and *zmiz1a* mutant embryos. A-B Ventral view of a 7 dpf embryo showing expression of *rag1* in the thymus. C-F. Lateral views of 4 dpf control and mutant fish. C-D. Visualization of green fluorescent neutrophils in the *Tg(mpx:eGFP)* $^{i114}$ fish. E-F. Visualization of macrophages in the *Tg(mpeg1:mCherry)* $^{gl22}$ fish.

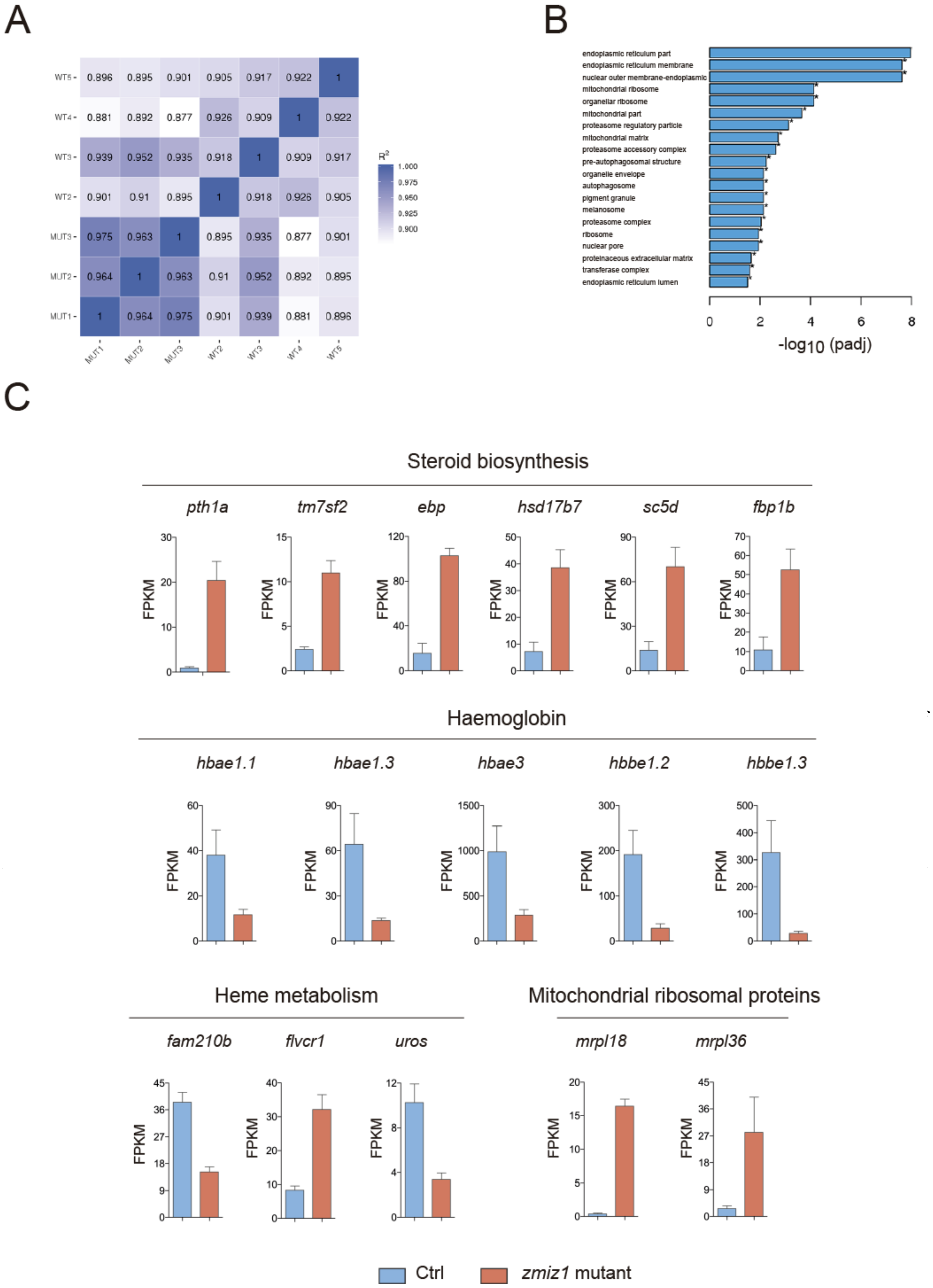
A
WT5
WT4
WT3
WT2
MUT3
MUT2
MUT1
R2
1.000
0.975
0.950
0.925
0.900
B
endoplasmic reticulum part
endoplasmic reticulum membrane
nuclear outer membrane-endoplasmic
mitochondrial ribosome
organellar ribosome
mitochondrial part
proteasome regulatory particle
mitochondrial matrix
proteasome accessory complex
pre-autophagosomal structure
organelle envelope
autophagosome
pigment granule
melanosome
proteasome complex
ribosome
nuclear pore
proteinaceous extracellular matrix
transferase complex
endoplasmic reticulum lumen
-log10 (padj)
C
Steroid biosynthesis
pth1a
tm7sf2
ebp
hsd17b7
sc5d
fbp1b
FPKM
Haemoglobin
hbae1.1
hbae1.3
hbae3
hbbe1.2
hbbe1.3
Heme metabolism
fam210b
flvcr1
uros
Mitochondrial ribosomal proteins
mrpl18
mrpl36
Ctrl
zmiz1 mutant

Supplementary Figure 4.

A. Pearson correlation coefficient matrix. $R^2$: Square of Pearson correlation coefficient (R). B. GO enrichment bar plot for cellular component (CC). The top 20 significantly enriched terms in the GO enrichment analysis. C. Bar charts comparing the normalized average expression levels (FPKM) for the indicated group of genes in control and *zmiz1a* mutant samples.
